# Conserved codon adaptation in highly expressed genes is associated with higher regularity in mRNA secondary structures

**DOI:** 10.1101/2020.11.23.393322

**Authors:** Mark G. Sterken, Ruud H.P. Wilbers, Pjotr Prins, Basten L. Snoek, George M. Giambasu, Erik Slootweg, Martijn H.M. Holterman, Johannes Helder, Jan E. Kammenga, Arjen Schots, Jaap Bakker, Lotte B. Westerhof

## Abstract

The redundancy of the genetic code allows for a regulatory layer to optimize protein synthesis by modulating translation and degradation of mRNAs. Patterns in synonymous codon usage in highly expressed genes have been studied in many species, but scarcely in conjunction with mRNA secondary structure. Here, we analyzed over 2,000 expression profiles covering a range of strains, treatments, and developmental stages of five model species (*Escherichia coli, Arabidopsis thaliana, Saccharomyces cerevisiae, Caenorhabditis elegans*, and *Mus musculus*). By comparative analyses of genes constitutively expressed at high and low levels, we revealed a conserved shift in codon usage and predicted mRNA secondary structures. Highly abundant transcripts and proteins, as well as high protein per transcript ratios, were consistently associated with less variable and shorter stretches of weak mRNA secondary structures (loops). Genome-wide recoding showed that codons with the highest relative increase in highly expressed genes, often C-ending and not necessarily the most frequent, enhanced formation of uniform loop sizes. Our results point at a general selective force contributing to the optimal expression of abundant proteins as less variable secondary structures promote regular ribosome trafficking with less detrimental collisions, thereby leading to an increase in mRNA stability and a higher translation efficiency.

## INTRODUCTION

Diverging frequencies of synonymous codons can be found in all domains of life and vary with the subsets of genes or gene regions compared. Ever since codon bias in highly expressed genes was shown to correlate with tRNA abundance profiles (optimal codons) (1–6), the general concept that codon use influences translational efficiency was adopted. More recent studies reveal that codon usage offers organisms an additional regulatory layer to govern a multitude of processes ranging from transcription, mRNA stability, translation initiation and elongation, to protein folding (7, 8).

An increasingly recognized mechanism to optimize the biogenesis of proteins is the optimization of translation speed along mRNA transcripts by adjusting codon use to subcellular tRNA concentrations (7, 8). Slowly translated codons at the beginning of mRNAs have been suggested to reduce ribosome traffic jams at later stages during elongation of abundant proteins (9, 10). Clustering of identical synonymous codons - irrespective their match with optimal codons - may enhance translation speed by recycling of nearby recharged tRNA molecules (11). Other forces shaping the codon landscape include the use of non-optimal codons to reduce the elongation rate to facilitate co-translational protein folding (12)and translocation of membrane proteins (13, 14). Alternatively, the kinetics of codon-anticodon pairing can also be tuned by modulating tRNA concentrations. Oscillating tRNA concentrations with anti-codons that enhance or repress translation speed have been implicated in tweaking up- and downregulation of gene expression during cell proliferation and differentiation.

An additional regulatory layer that controls the rate of translation is the secondary structure of mRNAs. Strong structures (low free energy) are demonstrated to slow down ribosomes and decrease the speed of translation (15–21). Generally, strong structures at the 5’ end of mRNAs are considered as disadvantageous for translation initiation (20, 22). Genome-wide analyses demonstrate well-conserved patterns across prokaryotes and eukaryotes towards reduced stability of mRNA structures at the initiation site, especially for GC-rich genes (23). However, weak folding of 5’ mRNA structures in *Escherichia coli* and *Saccharomyces cerevisiae* could not be correlated with expression levels of individual genes, and have been suggested to enhance the translation efficiency in a global manner (17). Another systematic trend in mRNA structures is described for the region directly downstream of the start codon in which strong mRNA structure is seen in *S. cerevisiae,* probably as a mechanism to slow down ribosome speed at the beginning and to avoid ribosome jamming during elongation (10, 24). In addition, it has been suggested that strong mRNA structures facilitate translational pauses to accommodate co-translational folding of compact proteins (25). Recently, it has been shown that certain mRNA regions appear to be subjected to conserved selective forces (26). Studying more than 500 organisms revealed in most phyla a strong folded region downstream of the start codon and weak secondary structures at the start and the end of coding regions (26).

Codon optimality can also have a profound effect on mRNA stability (8, 27, 28). Replacing optimal codons in budding yeast by less favorable codons significantly reduces mRNA stability, while reverse substitutions lead to a decreased decay (27). This positive correlation between codon optimality and mRNA stability is well-conserved and has been observed in *E. coli* (29)*, Xenopus* (30), zebrafish (30), as well as in mammalian cell lines (31). In *Saccharomyces cerevisiae* (32) the DEAD-box protein Dhh1p is required for the coupling between codon content and mRNA decay (32). Dhh1p is thought to sense the speed of ribosomes during elongation as it binds preferentially to transcripts enriched with slowly translated codons, thereby activating mRNA degradation. Together these findings indicate that codon usage may optimize gene expression via two distinct mechanisms that are linked via the dynamics of ribosome trafficking along transcripts: the kinetics of codon-anticodon pairing and mRNA turnover. The potential impact of ribosome speed on gene expression has been supported by analyzing codon choice in weak and strong mRNA secondary structures (33). In *E. coli* and *S. cerevisiae* optimal codons are preferentially used in regions with strong secondary structures, and codons matching with less abundant tRNAs are located in lowly structured regions (33). These opposing effects are considered as a mechanism to smoothen overall translation rates to optimize gene expression by reducing detrimental ribosome collisions (33).

Here we searched for patterns in codon usage and predicted mRNA secondary structures in constitutively highly expressed genes. Bias in codon usage has frequently been reported for highly expressed genes (7, 22, 34, 35), but scarcely in conjunction with structural features in mRNA folding. In addition, many studies (2, 3, 7, 36–38) focused on preferential codon usage in highly expressed genes and the match with cognate tRNA species (optimal codons). Here, we studied codon adaptation by analyzing relative changes in codon frequencies in highly expressed genes when compared with lowly expressed genes. This approach enables a comparative analysis of codon adaptation across species despite differences in optimal codons. By genome-wide comparisons of highly and lowly expressed genes, we found remarkably conserved patterns over five model species: *Escherichia coli, Arabidopsis thaliana, Saccharomyces cerevisiae, Caenorhabditis elegans,* and *Mus musculus.* In most species the relative changes in codon usage in highly expressed genes was associated with a reduction of mean and maximum loop sizes. This shift in mRNA secondary structures was found for highly abundant transcripts and proteins, as well as for high protein per transcript ratios. By recoding the genomes of the five model species, we show that codons with the highest relative increase in highly expressed genes reduces the variance and length of the loop sizes. Except for mouse, the overrepresentation of C-ending and avoidance of A- and T-ending codons in highly expressed genes was a commonly observed pattern across all species. Our findings suggest that the conserved bias in codon usage and associated smoothening of mRNA secondary structures reflect a general selection mechanism to enhance the expression level of abundant proteins, most likely via reducing detrimental ribosome collisions and promoting regular translation rates along transcripts.

## MATERIAL AND METHODS

### Data analysis

Data analysis was done in “R” (version 3.4.2, ×64) with custom written scripts (39). Data organization was mainly done using the tidyverse packages (40), and most plots were generated using ggplot2 (41). Scripts and the underlying transcriptome and structural database assembled to derive the figures and tables presented in the paper are accessible via a git repository: https://git.wur.nl/published_papers/sterken_codon_2020.

### Transcriptome collection, normalization, and transformation

Transcript abundance datasets of five species *(Escherichia coli, Arabidopsis thaliana, Saccharomyces cerevisiae, Caenorhabditis elegans,* and *Mus musculus*) were downloaded from Gene Expression Omnibus (GEO). Transcript abundance sets were selected based on abundant availability of data from one platform (Affimetrix), release date (not before 2008), publication linked to the GEO set and number of samples in the study. In total, 2067 transcript-abundance profiles were collected, representing eight to nine different studies per organism. An overview can be found in Supplementary table S1. The obtained data was normalized per sample based on the deposited intensities. For the comparison of highly and lowly expressed genes, the expression data was rank-normalized, ranking the expression as a percentage within each sample (0% being the lowest expressed gene, 100% being the highest expressed gene). For the analysis, the averages per probe over all samples within a species were used. For analysis of the transcript data, the log_2_ intensities were transformed to a z-score by applying the formula

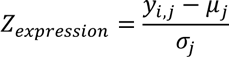

where the average (μ) log_2_ intensity over all spots of species *j* (j = 1,2, ..., 5) was subtracted from the log_2_ intensity (y) of spot *i* (varied per species) of species *j* and divided by the standard deviation (σ) over all the spots of species *j*. This assured that all data was comparable in range across species.

### Proteome collection and transformation

Protein abundances for each of the five species were obtained from the protein abundance database PaxDb (version 4) (42, 43). Abundances with a value of 0 were excluded from the set. As for the gene expression data, the protein abundance data was rank normalized for analysis of highest versus lowest protein abundances. Within each species the abundance was ranked as a percentage (0% the lowest and 100% the highest abundance).For analysis of the protein data, log_2_ transformed abundances were transformed to a z-score, as for the transcript abundance data.

### Protein per transcript ratio calculations

We collected both transcript abundance data and protein abundance data to calculate a protein per transcript ratio (PTR). This was calculated using the z-score data

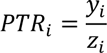

where PTR is the protein per transcript ratio of gene *i,* for which the log_2_ intensity transcript abundance (*y*) was divided by the log_2_ protein abundance *(z).* The PTR could only be calculated for genes where both transcript and protein abundance data was available.

### Coding sequences and mRNA structure prediction

The coding sequences (CDS) of all genes of five species were downloaded from sequence or genome repositories. For *Escherichia coli,* the CDS of strains CFT073, EDL933, MG1655 and Sakai were obtained from NCBI, accessions NC_004431.1, NC_002655.1, NC_U00096.3, and NC_002695.1 respectively. For *Arabidopsis thaliana,* the CDS of the 20101108 release were obtained from TAIR (44). For *Saccharomyces cerevisiae,* the open reading frames (without UTR, introns, etc.) of the 20110203 release were obtained from the Saccharomyces genome database (45). For *Caenorhabditis elegans*, the CDS of WS241 were obtained from WormBase (46). For *Mus musculus,* the CDS of the 20130508 release (GRCm38.p1) were obtained from the NCBI CCDS database (47).

The CDS datasets were filtered for mRNAs (by selecting coding sequences starting with ATG) and the mRNAs of all species were folded using Vienna RNA fold (version 2.1.8, x64 on Windows) (48), at 20°C, using the parameters of Andronescu, 2007 (49).

### mRNA sequence and structure statistics

Several statistics were taken from the mRNA sequence: gene length, absolute codon usage, absolute nucleotide usage (absolute number of A, T, C, and G) and the relative amount of each nucleotide per transcript, and nucleotide per codon position (e.g. the number of A at position 1), again absolute and relative.

The relative synonymous codon frequency (RSCF) for each codon was calculated across all selected transcripts using

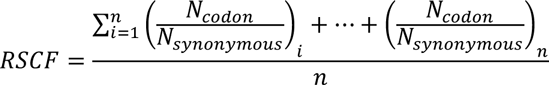

where *N_codon_* is the number of occurrences of a particular codon from the set of synonymous codons and *N_synonymous_* represents the number of occurrences of the synonymous codons, and *i* stands for a transcript of the *n* selected transcripts that contain the particular amino acid the synonymous codons encode. Thus, the values for RSCF vary between 0 and 1.

From the predicted mRNA structures ten parameters were taken: free energy (kcal/mol), number of bound nucleotides (nucleotides; nt), number of unbound nucleotides (nt), mean stem size (nt), mean loop size (nt), maximum loop size (nt), maximum stem size (nt), number of transitions from stem to loop (n), standard deviation of stem size (nt), standard deviation of loop size (nt) of the structure. For some of the factors a correction was applied for gene-length, these were: energy, number of bound nucleotides, number of unbound nucleotides, and number of transitions from stem to loop.

### Relative synonymous codon frequencies linked to mRNA abundance, protein abundance, and protein per transcript

Changes in frequencies for each codon were calculated using

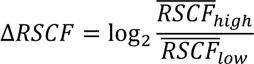

where the relative change in codon frequency (ΔRSCF) was calculated for the average codon frequency in the 5% highest expressed genes (RSCF_high_) versus the 5% lowest expressed genes (RSCF_low_). This resulted in a measure for changes in codon usage linked to expression levels. This method was applied for transcript abundances (relative synonymous codon use transcript abundance; ΔRSCF-TA), protein abundances (relative synonymous codon use protein abundance; ΔRSCF-PA), and protein per transcript ratios (relative synonymous codon use protein per transcript ratio; ΔRSCF-PTR). Comparison of the different ΔRSCF values within species was done using Pearson correlation as to account for ΔRSCF amplitude differences. To compare the direction of ΔRSCF associations between species, Spearman correlations were used. Spearman was used as the amplitudes of the ΔRSCF between species differed strongly.

### mRNA structures linked to mRNA abundance, protein abundance, and protein per transcript ratio

To evaluate the predicted mRNA structures in a similar way as shifts in codon frequencies, the relationship between mRNA structure and expression was calculated using

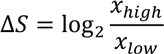

where the structural bias (ΔS) was calculated for the average structure value in the 5% highest expressed genes (x_high_) versus 5% lowest expressed genes (x_low_). This method was applied for transcript abundances, protein abundances, and protein per transcript ratio. Comparison of the different structural biases within species was done using Pearson correlation as to account for ΔRSCF amplitude differences.

### Permutation analyses of codon biases and mRNA structures

To assign probabilities to the observed trends, a permutation approach was used. Within each species the synonymous codons were randomly distributed over the amino acids. This resulted in an mRNA pool that encoded the same proteins and contained the same overall codon use. This procedure was conducted 100 times for each species, after which the permutated mRNA pool was folded and scored according to the procedures described above.

The resulting distributions were used to determine whether the trends in codon bias linked to TA, PA, and PTR were likely to be random or the result of a non-random process. We ranked the relative synonymous codon use values from each permutation, the 2^nd^ highest and 2^nd^ lowest were taken as the 95% confidence interval. Based on these values the significances of the codon biases were assessed. Like the trends in codon bias, these distributions were also used to determine if mRNA structure properties linked to TA, PA, and PTR were likely to be random or the result of a non-random process.

### Structure evaluation through mRNA sequence recoding based on preferred codons

To evaluate the impact of changes in codon usage on mRNA structure, the entire coding sequence library of each species was recoded by replacing each codon with the synonymous codon that showed the highest relative change (ΔRSCF) upon comparison of highly and lowly expressed genes. This recoding process was conducted for codons that were selected by analysing transcript abundances, protein abundances as wells protein per transcript ratios. A comparison between the native and recoded mRNAs was made per transcript by

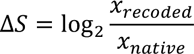

where the structural change (ΔS) was calculated for the average structure value in recoded genes (xrecoded) versus native encoded genes (x_native_). The ΔRSCF-identified codons used for recoding that showed the highest relative changes can be found in Supplementary Table S5. It is noted that these codons are not necessarily the most frequent synonymous codons. To test if recoded mRNA structures differed from the native encoded mRNA structures, a paired t-test was conducted on the untransformed structure features. In total, 11 structure features over five species with three different recodings were tested: codons that were selected by analysing ΔRSCF-TA, ΔRSCF-PA, and ΔRSCF-PTR data. A Bonferroni correction was applied to correct for multiple testing.

## RESULTS

### Conserved adaptation in codon usage in highly abundant transcripts

To determine whether global patterns in codon change can be identified in abundant mRNAs, we analysed for each of five model species at least 250 transcript profiles (Supplementary Table S1). After rank-normalization, the expression levels across developmental stages, tissues, strains and treatments were averaged to determine the relative ranking of the genes within each species. Next, the highest and lowest 5% expressed transcripts were selected and the relative synonymous codon frequency was determined per transcript (Figure 1A) and used to calculate to average relative codon frequency (RSCF) within the two sets of transcripts. To evaluate the relative change in codon usage between the two classes of transcript abundances (TA), the log_2_ ratio of the two RSCF values was taken as a measure (ΔRSCF-TA) that indicated if a particular codon occurred more (>0) or less (<0) frequently in genes with high transcript abundance as compared to genes with low transcript abundance (Figure 1B) (for further details see material and methods).

**Figure 1.**
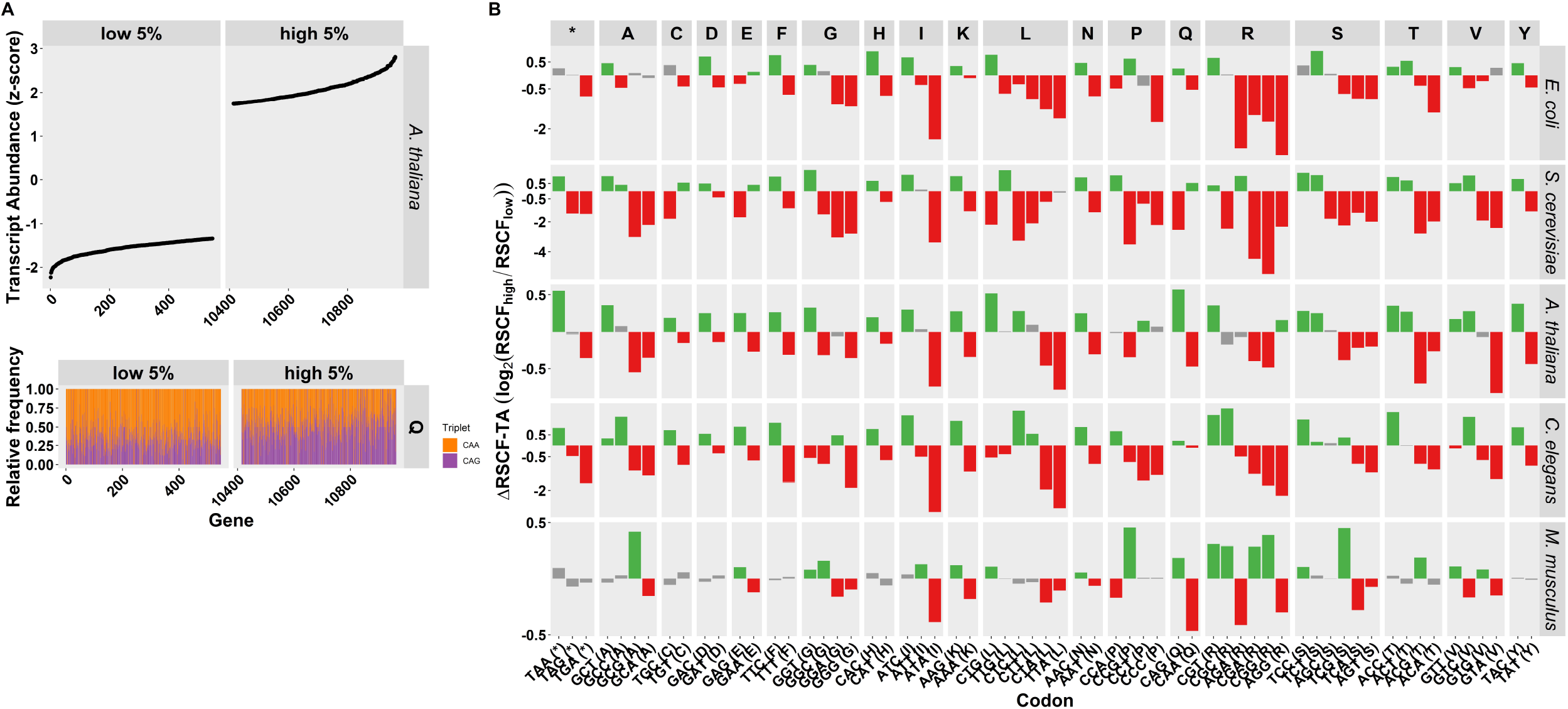
Relative change in codon frequency (ΔRSCF-TA) associated with transcript abundance. (A) Transcript abundance profiles were gathered from five model species. The genes of each species were ranked according to transcript abundance (upper panel - example *A. thaliana)* and the RSCF values were determined for the 5% highest expressed genes (RSCF_high_) and the 5% lowest expressed genes (RSCF_low_) (lower panel - example codons CAA and CAG encoding glutamine (Q)). (B) Shifts in codon frequencies (ΔRSCF-TA) calculated as the log_2_ ratio of RSCF values of the highest abundant 5% (RSCF_high_) and the lowest abundant 5% (RSCF_low_). Colours indicate if a particular codon is significantly positively associated (green) or negatively associated (red) with high transcript abundances (FDR < 0.05), grey bars indicate non-significant associations (FDR > 0.05).

The comparison of codon usage in highly and lowly expressed RNAs uncovered a remarkable similar pattern of over- and underrepresentation of codons across the evaluated species (Figure 1B). Yet, the amplitudes of the ΔRSCF-TA values varied strongly across species, where *E. coli* and *S. cerevisiae* displayed the largest and *M. musculus* the lowest amplitudes (Figure 1B, note different scales on Y-axes). *M. musculus* exhibited not only the smallest amplitudes, but also the least overlap in codon bias with the other four species. In *E. coli, S. cerevisiae, A. thaliana,* and *C. elegans* 27 codons (out of 62; excluding the codons for M and W) were significantly over- or underrepresented (permutation, FDR < 0.05) (Supplementary Table S2). These triplets encode 14 different amino acids (out of 18) and the termination of translation. Codons showing a positive association with high mRNA levels were often C-ending (9/19), whereas codons showing a negative relationship with high transcript abundances were mostly A- or T-ending (16/19). Apart from *M. musculus,* the direction of ΔRSCF-TA values correlated remarkably well over four species (Spearman correlation, R^2^ ranging from 0.41-0.71; Supplementary Figure S1), demonstrating that the codon biases associated with high transcript abundances were well-conserved.

### Transcript and protein abundances, as well as protein per transcript ratios reveal similar patterns in codon adaptation across species

To evaluate the codon usage in relation to protein levels, we retrieved data of the five species from the protein abundance database PaxDb (42, 43). The same analysis as for the transcript abundance was conducted, resulting in a measure of synonymous codon use related to protein abundance (ΔRSCF-PA), which indicate whether a codon was used more (>0) or less (<0) frequently in genes encoding highly abundant proteins (top 5%) than in genes with low abundances (lowest 5%) (Supplementary table S2; Supplementary Figure S2). Except for *M. musculus,* the ΔRSCF-TA and -PA values revealed a similar set of codons that were over- or underrepresented in genes encoding highly abundant proteins. Within the four species the ΔRSCF-TA and -PA values were strongly correlated with a Pearson R^2^> 0.84 (Supplementary Figure S3A). The relation between codon usage and protein levels was relatively well conserved between *E. coli, S. cerevisiae, A. thaliana,* and *C. elegans* as indicated by the correlations between the ΔRSCF-PA values (Spearman R^2^ between 0.46-0.74; Supplementary Figure 3B). Altogether, except for *M. musculus,* our results demonstrated that the shift in codon usage related to transcript and protein levels was highly similar within each species and was relatively well-conserved across species.

Next, we studied the relationship between codon usage and the amount of protein per transcript (Figure 2; Supplementary Table S2; Supplementary Figure S4). The protein per transcript ratios were rank-normalized, and the highest and lowest 5% values were selected to calculate the change in relative synonymous use (ΔRSCF-PTR) between the extremes (Supplementary Figure S5). Remarkably, the ΔRSCF-PTR values within species identified nearly the same codons as the ΔRSCF-TA values (except for *M. musculus;* Pearson R^2^ between 0.81 and 0.99; Figure 2B). Also, the correlation between ΔRSCF-PTR and ΔRSCF-PA values was very high within the four species with Pearson R^2^ between 0.95 and 0.99. It was noted that these high correlations were observed even though the underlying gene pools were distinct. The gene pools encoding lowly expressed transcripts and proteins and had low amounts of protein per transcript (Figure 2A) shared on average across four species only 4% overlap (Figure 2C). Similarly, pools analysed for the top 5% shared 20% of their genes (Figure 2D; Supplementary Figure S4 for absolute numbers). These data demonstrated that genes encoding highly abundant transcripts and proteins, as well genes having high protein/transcript values, selected similar codon-pools. In addition, apart from *M. musculus,* the pattern of codon bias exhibited a relatively high degree of conservation across species and was dominated by an increase in C-ending codons in highly expressed genes (Supplementary Figure S3B).

**Figure 2.**
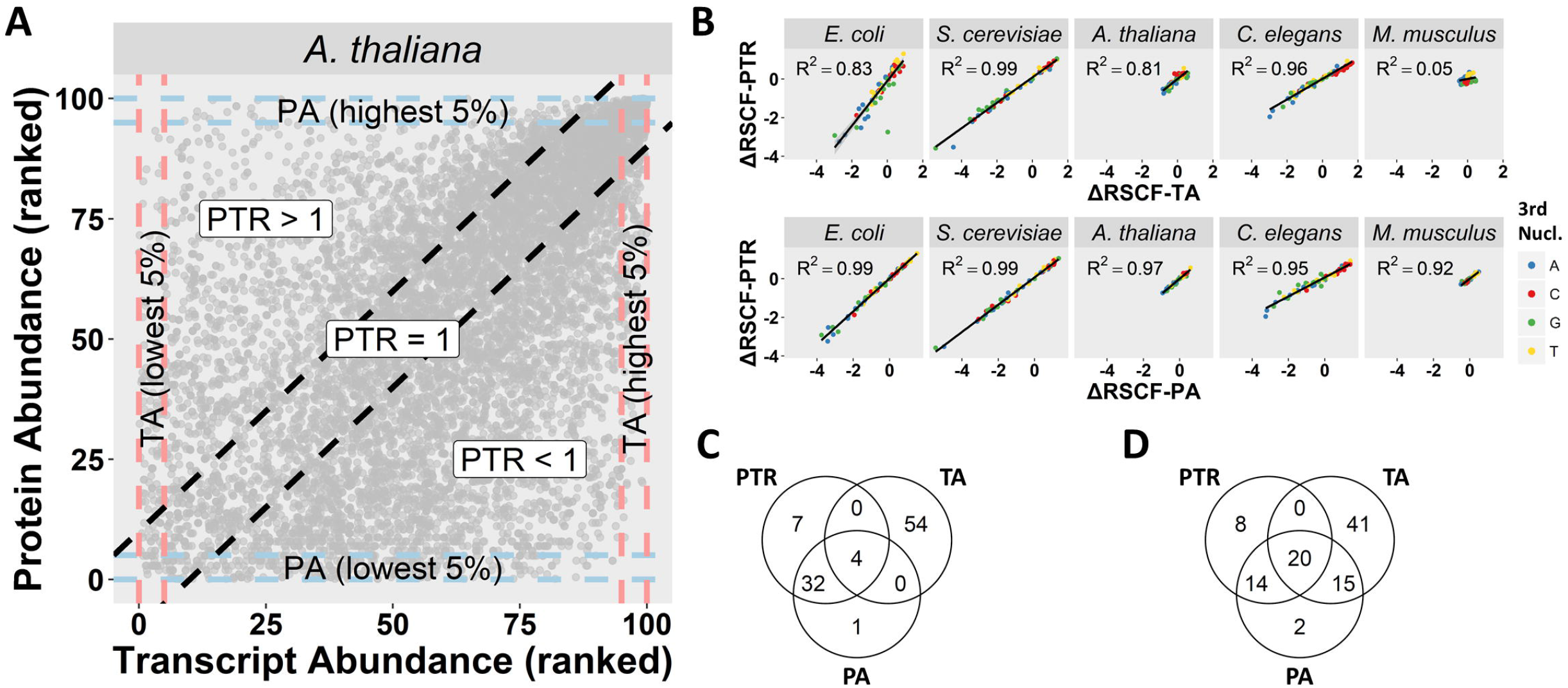
Relation between adaptation of codon usage associated with transcript abundance (ΔRSCF-TA) and protein abundance (ΔRSCF-PA) versus protein per transcript ratios (ΔRSCF-PTR). (A) A schematic overview of the relation between transcript abundance (TA) and protein abundance (PA), each dot shown represents a gene for which both values are known. The dashed red lines represent the areas of 5% lowest (left) and 5% highest (right) transcript abundances (TA). The dashed blue lines represent the areas of 5% lowest (bottom) and 5% highest (top) protein abundances (PA). The dashed black lines are a visual marker to indicate where protein per transcript ratio (PTR) equals 1, mind that this is only a visual presentation, the actual PTR values are calculated using z-transformed expression data. (B) The correlations between ΔRSCF values calculated for TA and PA versus PTR. Each dot represents a triplet, the colour indicates the letter of the 3^rd^ nucleotide. The R^2^ reported is calculated from the Pearson correlation-coefficient. (C) Venn-diagram of the percentage overlap in genes comprising the lowest 5% of TA, PA, and PTR. The values are averaged over all species except mice. (D) As in C, but for the genes making up the highest 5% of TA, PA, and PTR.

### Protein and transcript abundances and their ratios are positively associated with uniform mRNA secondary structures

To study whether the patterns in codon usage across species was reflected in general characteristics of mRNA secondary structures, we decided to compare the predicted mRNA structures using Vienna RNA fold (48). The methodology to evaluate the predicted structural characteristics of mRNAs was similar to the analyses of codon usage. The bias in secondary structures was determined by calculating the log_2_ ratio for each structural feature (ΔS), which indicated whether a particular characteristic was encountered more (>0) or less frequently (<0) in highly (top 5%) than in lowly (bottom 5%) expressed transcripts (Figure 3C, Supplementary Table S3). Comparison of the extremes showed that transcript abundance was positively linked to more stable mRNA structures (lower free energy), number of transitions from stem to loop, and the number of bound nucleotides (permutation, FDR < 0.05). A negative linkage was found between transcript abundance and the mean number of unbound nucleotides in loops (loop size), standard deviation of loop size, and maximum loop size (permutation, FDR < 0.05). Overall, it can be said that highly abundant transcripts are on average more uniform in the sense that they have smaller and less variable loop sizes (Figure 3C, Supplementary Table S3).

**Figure 3.**
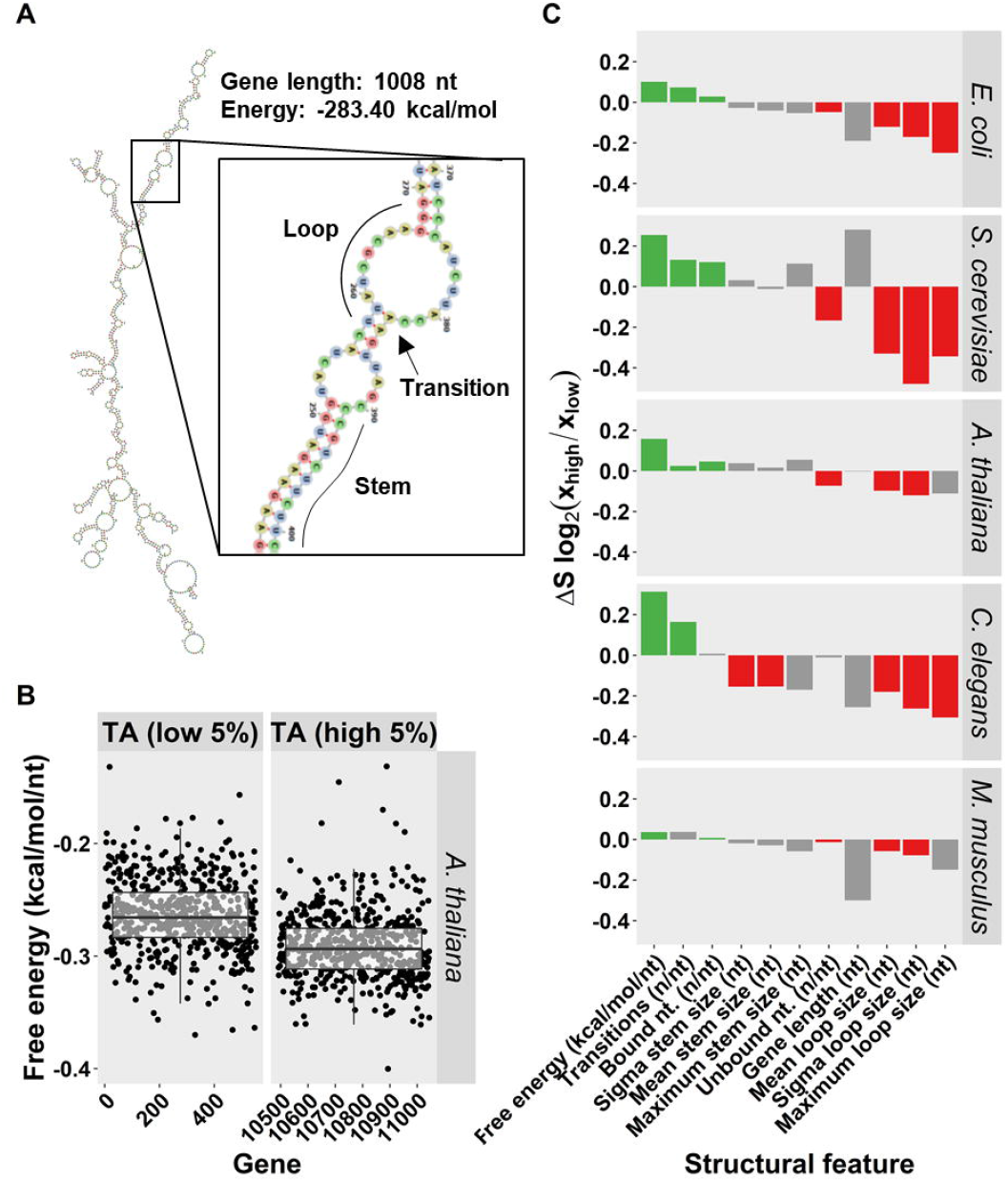
Bias in structural features of mRNA folding associated with transcript abundance (TA). (A) A predicted mRNA secondary structure (of the BZR1 gene of *A. thaliana),* modelled using RNAfold and visualized using forna (75). In the enlargement, three structural features that were quantified are indicated: a stem, a loop, and a transition. (B) The contrast in free energy (kcal/mol/nt) between the 5% lowest abundant transcripts and the 5% highest abundant transcript in *A. thaliana.* Each dot represents a gene. The boxplot is added for visual purposes. (C) Bias in structural features (ΔS) calculated as the log_2_ ratio of the average structure value (e.g. number of bound nucleotides (nt)) in the highest abundant 5% (x_high_) and the lowest abundant 5% (x_low_). Colours indicate if a particular feature is significantly positively associated (green) or negatively associated (red) with high transcript abundances (FDR < 0.05), grey bars indicate non-significant associations (FDR > 0.05).

A higher degree of uniformity in mRNA secondary structures was also found for high protein abundances and, to lesser extent, for high protein per transcript ratios (Supplementary Figure S6, Supplementary Table S3). At high protein abundances, the maximum loop sizes were significantly smaller in all species and, apart from *M. musculus,* accompanied with a reduction in average length of the loops and lower standard deviations in loop lengths. For both PA and PTR the number of stem-loop transitions was positively related to high protein abundances or high protein per transcript ratios for *E. coli, S. cerevisiae, A. thaliana,* and *C. elegans* (permutation, FDR < 0.05). For the same four species, a reduction in mean loop sizes and lower standard deviations in loop lengths were observed with increasing amounts of protein per transcript (permutation, FDR < 0.05). Despite the increase in G/C content and reduction of unbound nucleotides at high PA and PTR extremes, only in a few cases a significant increase of stronger structures (lower free energy) could be noticed (Supplementary Figure S6, Supplementary Table S3).

In conclusion, despite the fact that largely non-overlapping gene pools were sampled (Figure 2C and 2D, Supplementary Figure S4), all three types of analyses (TA, PA and PTR) revealed a similar pattern. At high extremes all species, although to a lesser degree for *M. musculus,* showed a significant bias towards more uniform loop sizes, i.e., a reduction of mean, maximum and standard deviation of stem size(s) (Figure 3C, Supplementary Figure S6, Supplementary Table S3). Another frequently observed pattern at high extremes was a rise in the n5umber of stem-loop transitions (Figure 3C, Supplementary Figure S6). Altogether, our data indicated that highly expressed genes tend to have more regular mRNA secondary structures, predominantly mediated by smaller and less variable loop sizes.

### Codons with highest relative increase in highly expressed genes enhance formation of uniform mRNA secondary structures

To determine the influence of the conserved codon adaptation on secondary mRNA structures, the genomes of the five model species were recoded by selecting for each amino acid a codon that increased most in frequency at high extremes as identified by the TA, PA, or PTR analyses. It should be noted that these selected codons were not necessarily the most frequently used codons in highly expressed genes. In several cases, codons with the highest frequency in highly expressed genes did not show the highest relative increase. Depending on the species and the type of analysis, in three to seventeen cases the most frequent codon was not the same as the codon with highest value for ΔRSCF (Supplementary Figure S7).

For most amino acids the selected codons largely overlapped between the three types of analyses, not only within species but also between species (Supplementary Table S4). After recoding, the secondary structures were predicted, and the structural characteristics of each recoded mRNA was compared with those of its native structure. This comparison revealed that codons showing the highest relative increase at high transcript abundances (TA) caused a significant bias in secondary structures by increasing the physical stability of the secondary mRNA structure, the number of stem to loop transitions, and the number of bound nucleotides (Bonferroni adjusted paired t-test, p < 0.05; Figure 4). Furthermore, a decrease in maximum loop size, standard deviation of the loop size, mean loop size, and number of unbound nucleotides was observed (Figure 4; Supplementary Table S5). Similar patterns were caused by codons showing the most positive relationship with high protein abundances (PA) and high protein per transcript ratios (PTR) (Supplementary Figure S8; Supplementary Table S5).

**Figure 4.**
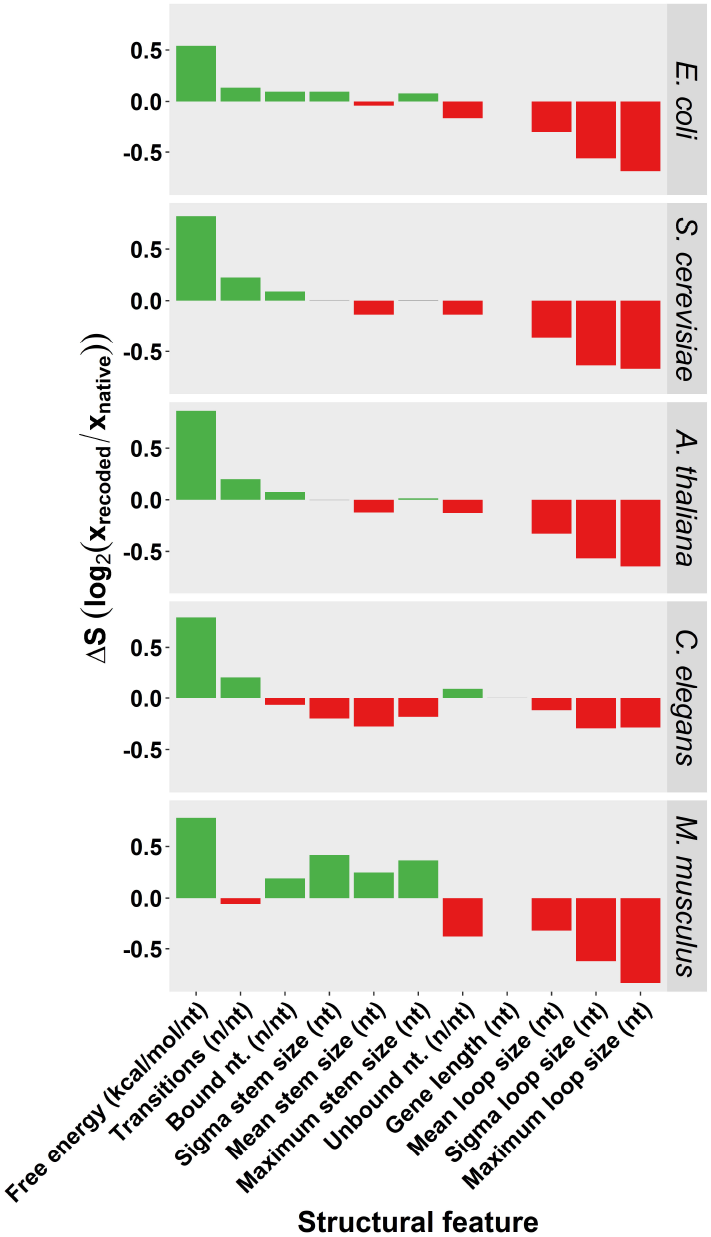
Changes in mRNA secondary structures after recoding with codons that showed the highest relative increase in highly expressed transcripts. Each recoded transcript was compared to its native counterpart. The shifts in structural features (ΔS) were calculated as the log_2_ ratio of the average structure value (e.g. number of nucleotides (nt) per maximum loop size) in recoded sequences (xrecoded) and in native sequences (x_native_). Only the sequences for which transcription data was assessed were evaluated to make results comparable (n = 2,782 for *E. coli,* n = 5,655 for *S. cerevisiae,* n = 13,569 for *A. thaliana,* n = 17,657 for *C. elegans,* and n = 16,938 for *M. musculus).* Colours indicate if a particular feature is significantly positively associated (green) or negatively associated (red) with high transcript abundances (Bonferroni corrected paired t-test, p < 0.05), grey bars indicate non-significant associations (Bonferroni corrected paired t-test, p > 0.05).

In conclusion, recoding the genes of the five model species demonstrated that codons with the largest relative frequency changes at high TA, PA and PTR levels enhanced the formation of regular secondary structures, mainly through reducing the size and variation of the loops.

### Interdependence between mRNA structural features

To obtain insight in the interdependence between the mRNA structural features analysed in this study, we examined the correlations between mRNA characteristics in native and recoded mRNAs in the five model species (Supplementary Figure S9 and S10). As expected, both mean and standard deviations of loop sizes were consistently negatively correlated with maximum loop sizes. Similar correlations were observed for mean, maximum and standard deviation of stem size(s). Other mRNA structural features that were intertwined and consistently, either negatively or positively, correlated across species were: average loop sizes versus number of bound nucleotides, free energy versus number of (un)bound nucleotides (further details see Supplementary Figure S9 and S10). No correlations were observed between mean, maximum and standard deviation (σ) of loop and stem size(s). on the one hand, and – on the other – mean, maximum and standard deviation (σ) of stem size(s). This independence between loop size and stem size is also reflected in Figure 4, where the mean, sigma and maximum loop sizes in *C. elegans* were negatively linked with mean, sigma and maximum stem sizes, whereas in *M. musculus* a reverse relationship was observed.

## DISCUSSION

This work presents a comparative analysis of the most highly and lowly expressed genes in more than 2,000 expression profile and reveals a conserved shift in codon usage and predicted mRNA secondary structures. Significances were determined through permutation of codon use, where the synonymous codons were randomly distributed for each gene while the overall codon use and protein products were kept similar. This approach allowed us to define whether the codon-bias was non-random and not linked to other factors that could affect our analyses, including amino acid composition and gene length. The change in codon usage in highly expressed genes, characterized by an increase in C-ending codons and avoidance of A- and T-ending codons, is associated with a reduction of the mean and maximum loop sizes and an increase in the number of stem-loop transitions. These patterns are observed for high transcript and protein abundances and, to lesser extent, also for high protein per transcript ratios. Altogether our data point at a general selective force that optimizes both mRNA stability and translation efficiency.

Genome-wide recoding of mRNAs with codons that showed the largest relative changes in frequencies in highly expressed genes resulted in more uniform secondary structures in all five model species. Compared to native sequences, the recoded sequences promoted a consistent reduction in mean, standard deviation and maximum loop size and an increase of stronger structures. Stem sizes were less affected by recoding and the effect varied across species. Altogether, this suggests that the conserved codon bias at high extremes of TA, PA, and PTR, predominantly influence loop size features. The most likely explanation is that the shift towards C-ending codons and avoidance of A and T-ending codons enhances the propensity to form stable secondary structures, which reduces loop size but apparently does not necessarily result in an increase in stem size.

Codon bias across multiple organisms has been well documented in literature. Already in the 1980s studies reported on tRNA abundances correlating with codon usage (5, 50). Codon adaptation has been linked to both transcript and protein abundance (8, 22). Genome-wide codon bias is driven by weak selective forces, mutation and genetic drift (6). While selection has been demonstrated to play a dominant role of in non-mammalian species (36, 51), in mammalian species mutational biases are more prominent (52, 53) and the contribution of selection is more disputed (6, 22). This correlates well with our findings where the conserved bias in codon use and mRNA secondary structures in *M. musculus* is less pronounced than in the other four model species. The approach used in this work identifies codon adaptation by calculating the ratio of the relative coding frequencies (RSCF) in highly and lowly expressed genes. Our computational method minimizes the contribution of mutation bias (54) and focuses on the effect of selection, and is conceptually similar to that presented in (54), although it does not distinguish between A/G-ending or T/C-ending codons. Other methods to identify codon-biases exist, including codon adaptation index (CAI) (37), translation adaptation index (tAI) (55), and codon deviation coefficient (CDC) (56). All methods have their interpretations and biases, which are still topic of debate (54). A key advantage of the current approach is that it simply compares the relative enrichment of the use of a specific synonymous codon in a group of genes defined by a measurement of abundance. The pronounced outcome of our analyses may be attributed to evolutionary forces that act both on highly and lowly expressed genes. While traditionally selective forces have been proposed to be the most dominant at high expression levels (6), recent work has suggested that codon usage in lowly expressed genes is subjected to selection and reduces the stability of the mRNA secondary structures (57).

Codon optimality, and hence translation rate, is thought to promote mRNA stability via regulating the rate of elongation (8, 58). The positive relationship between translation rate and mRNA stability has been reported for a diverse range of organisms ranging from *E. coli* (29), *Drosophila* (59), zebrafish (30), *Xenopus* (30), to humans (60, 61). In *S. cerevisiae* the DEAD-box protein *Dhh1p* has been shown to monitor ribosome densities and to target inefficiently translated mRNAs for degradation (32). In view of these observations, we assume that the observed bias in mRNA secondary structures contributes to the stability of highly abundant transcripts by reducing variation in ribosome speed along transcripts. The translation rate in weakly structured regions (loops) is relatively high and may lead to congestion of ribosomes at the start of highly structured regions, thereby leading to collisions and enhanced mRNA decay. An alternative explanation, not mutually exclusive, is that larger loop sizes with inherent higher ribosome speed have larger stretches of ‘naked’ mRNA and are less protected by ribosomes that shield mRNAs from degradation by endonucleases (62). Since codon usage and mRNA secondary structures are intertwined, smoothening of the elongation rate is expected to be a delicate interplay between optimizing codon composition and mRNA folding. The existence of such an interplay has been reported for *E. coli* and *S. cerevisiae* where a reduction of fluctuating translation rates is achieved by an increase in optimal codons in highly structured regions (stems), and a preferential use of non-optimal codons in weakly structured regions (loops) (33).

The notion that the avoidance of local fluctuations in ribosome densities enhances mRNA stability may explain that both high transcript abundances and high protein per transcript ratios are associated with a similar bias in codon usage and inherent mRNA secondary structures. Local fluctuations in translation rates leading to ribosome collisions and stalling may trigger no-go decay of the transcript and degradation of the partly synthesised protein (63–68). Modelling studies (69) emphasize that varying rates of translation may be unproductive and indicate that evenly spaced distances between ribosomes are more beneficial for optimal elongation rates. Thus, the bias in mRNA secondary structures associated with a high protein per transcript ratio indicates that smoothening of secondary structures may enhance translation efficiencies. It is stressed that the amount of protein per transcript seems a fair proxy for translation efficiency, as genome-wide protein turnover plays a rather small role in the explanation of protein abundance variation (70, 71). Remarkably, while smoothening of the secondary structures is positively linked with transcript levels (TA) as well as protein per transcript ratios (PTR), the underlying gene pools hardly overlap. These data imply that our findings are independent of the function and sequence structure of the underlying genes indicating that two different pathways converge by selecting codons and inherent mRNA structures that promote regular translation rates along transcripts. It is noted that protein abundances (PA) are the output of transcript levels (TA) and the number of polypeptides synthesized per mRNA (PTR) and, consequently, reveal a similar bias at high extremes as the TA and PTR analyses, which is also reflected in the large overlap of the underlying genes at high expression levels.

In sum, we suggest that the high correlations between all three types of analyses (TA, PA and PTR) can be explained by selection towards mRNA structures that facilitate a uniform ribosome speed during elongation. However, the strength and relevance of the underlying mechanism remains elusive. A multitude of selective forces (7, 8, 20–22, 72, 73) optimizes the biogenesis of transcripts and proteins and various regulatory mechanisms may have opposing roles in selecting particular codons and, hence in shaping mRNAs structures, and may weaken the observed bias in secondary structures. For example, reducing fluctuations in ribosome trafficking may also be accomplished by other processes, like selecting optimal codons in highly structured regions and less beneficial codons in lowly structured regions (33). Evidently, we do not exclude other mechanisms that may link the conserved bias revealed by all three types of analyses. Further large-scale studies are required to unravel the underlying mechanism(s). For example, by testing synthetic genes with mRNA structures varying in loop and stem sizes while keeping factors like overall codon usage and thermodynamic properties constant in combination with ribosome profiling and mRNA stability studies.

Irrespective the underlying mechanisms, our results provide an additional guideline in codon optimization strategies for the design of synthetic genes. Although many successes have been reported, results are unpredictable and optimizing codon use remains a challenge (74). To increase the production of proteins in heterologous expression systems, several methods have been developed that optimize expression by adapting the CAI’s index to the expression host, together with adjusting sequence features that often include GC content, avoidance of Shine-Dalgarno-like sequences, transcriptional terminator sites and RNase recognition sites (74). Our findings can be used to extend existing algorithms to design synthetic genes leading to higher expression levels by selecting codons that lead to more uniform mRNA secondary structures.

Altogether, our analysis across four kingdoms of life reveals a broad selection pressure towards more uniform mRNA structures. Genome-wide recoding of the mRNAs with codons that are increase most in highly expressed genes results in a shift towards smaller and less variable mRNA loop sizes. Our data indicate that selection on regular mRNA secondary structures may contribute to shaping the codon landscape and plays next to codon-anticodon adaptation and various other mechanisms (7, 8, 72, 73), a significant role in enhancing the synthesis of highly abundant proteins. A plausible mechanism to explain our observations is that more regular mRNA structures promote gene expression by reducing detrimental fluctuations in ribosome trafficking that may attenuate both mRNA stability and translation efficiency.

## AVAILABILITY

Scripts and the underlying transcriptome and structural database assembled to derive the figures and tables presented in the paper are accessible via a git repository: https://git.wur.nl/published_papers/sterken_codon_2020.

## ACCESSION NUMBERS

Accession numbers of the transcriptome studies used can be found in Supplementary Table S1.

## SUPPLEMENTARY DATA

Supplementary Data are available along with the publication.

## ACKNOWLEDGEMENT

We want to thank Aska Goverse and Geert Smant for their comments on early versions of this manuscript.

## AUTHOR CONTRIBUTIONS

R.H.P.W., P.P., A.S. and L.B.W. conceived the idea at the basis of this manuscript. J.H., L.B.S., J.E.K., A.S., J.B. assisted with the design of the analyses and the interpretation. M.G.S., L.B.W. and J.B. wrote the manuscript with input from G.M.G., J.H., A.S. and J.E.K.. M.G.S. analysed and visualized the data with input from P.P, L.B.S., and M.H.M.H.. G.M.G. assisted with the interpretation of mRNA folding data. M.H.M.H. collected and curated the transcriptomics data. L.B.S. normalized the transcriptomics data. P.P. conducted a pilot analysis on ΔRSCF values and secondary mRNA structures. All authors commented on draft versions of the manuscript.

