## Supplementary legends and figures for "Conserved codon adaptation in highly expressed genes is associated with higher regularity in mRNA secondary structures"

**Supplementary Table S1**. An overview of the microarray studies used to determine transcript abundances. In the first tab an overview is provided of the number of samples per species. The second till sixth tab gives a detailed overview per species (*E. coli*, *A. thaliana*, *S. cerevisiae*, *C. elegans*, and *M. musculus*).

*Table is provided as an .xlsx file*

**Supplementary Table S2.** The relative change in codon frequency (ΔRSCF) based on transcript abundance (ΔRSCF-TA), proteins abundance (ΔRSCF-PA), and protein per transcript ratio (ΔRSCF-PTR) for all five species and all codons. Only for ATG (M) and TGG (W) no values could be calculated as these amino acids are not encoded by redundant codons. The asterisk (*) indicates if the association is significant (permutation, FDR < 0.05), whereas (NS) indicates if this is not the case.

*Table is provided as an .xlsx file*

**Supplementary Table S3.** The differences in mRNA structure between top and bottom 5% for transcript abundance, proteins abundance, and protein per transcript ratio for all five species**.** The following structural characteristics are listed: free energy per nucleotide (kcal/mol), fraction of bound nucleotides, fraction of unbound nucleotides, mean stem size (nt), mean loop size (nt), maximum loop size (nt), maximum stem size (nt), relative number of transitions from stem to loop (n/nt), standard deviation of stem size (nt), standard deviation of loop size (nt), and gene length (nt). The asterisk (*) indicates if the association is significant (permutation, FDR < 0.05), whereas (NS) indicates if this is not the case.

*Table is provided as an .xlsx file*

**Supplementary Table S4.** A matrix with the codons used for recoding the entire transcriptome for the five species. The codons were chosen by selecting for each amino acid a codon that increased most in frequency at high extremes as identified by analyses of transcript abundances (ΔRSCF-TA), protein abundances (ΔRSCF-PA) and protein per transcript ratio (ΔRSCF-PTR). The codons are organized per species and per type (ΔRSCF -TA, ΔRSCF -PA or ΔRSCF -PTR). For most amino acids only one codon was the most strongly associated with high expression, but there were four codons on a shared first place. Three of these were found in mice for ΔRSCF -PTR, and one was found for the stop-codon for ΔRSCF-PTR in Arabidopsis.

*Table is provided as an .xlsx file*

**Supplementary Table S5**. Effect of whole-genome recoding on mRNA secondary structures. The values indicate the relative changes in structural features (ΔS) and were calculated as the $\log_2$ of the ratio for the average structure value (e.g. number of nucleotide per mean stem size) in recoded genes ($x_{recoded}$) versus native encoded genes ($x_{native}$). The recoding was carried out for three sets of selected codons based on the analyses of transcript abundances (TA), protein abundances (PA) and protein per transcript ratios (PTR). The impact of recoding on structure parameters are given per species and for each recoding procedure. Asterisk (*) indicate the significance according to a Bonferroni-corrected paired t-test ($p < 0.05$). In case there was no significant difference, an NS was added.

*Table is provided as an .xlsx file*

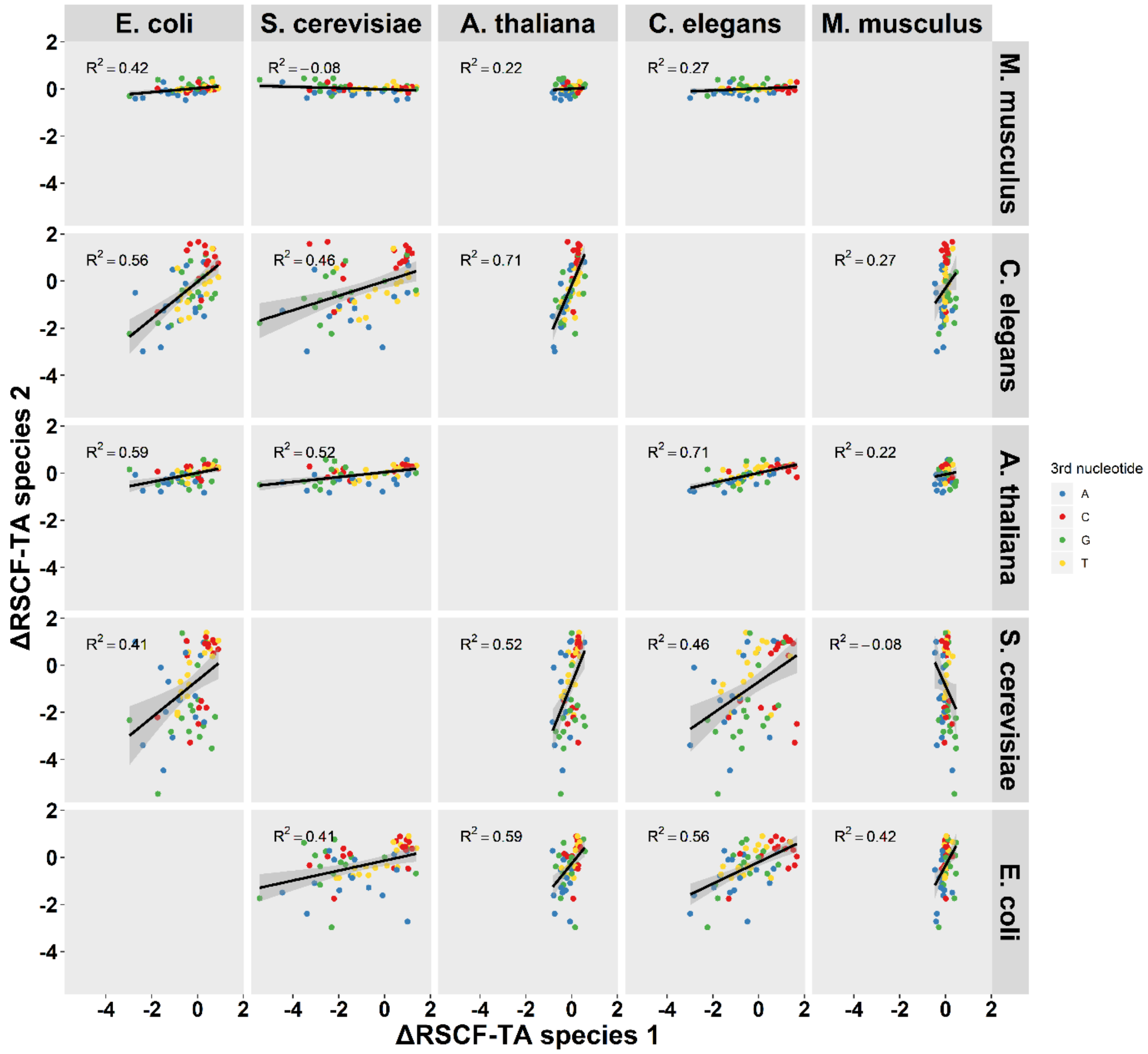

**Supplementary Figure S1.** The relation between shifts in codon frequencies associated with transcript abundance (ΔRSCF-TA) between species. On the x-axis the ΔRSCF-TA values for species 1 are plotted (species name is indicated on the bar on top), on the y-axis the ΔRSCF-TA values for species 2 are plotted (species name is indicated on the bar on the right). The line is based on a linear model and added as visual aid. The reported $R^2$ is based on the spearman correlation.

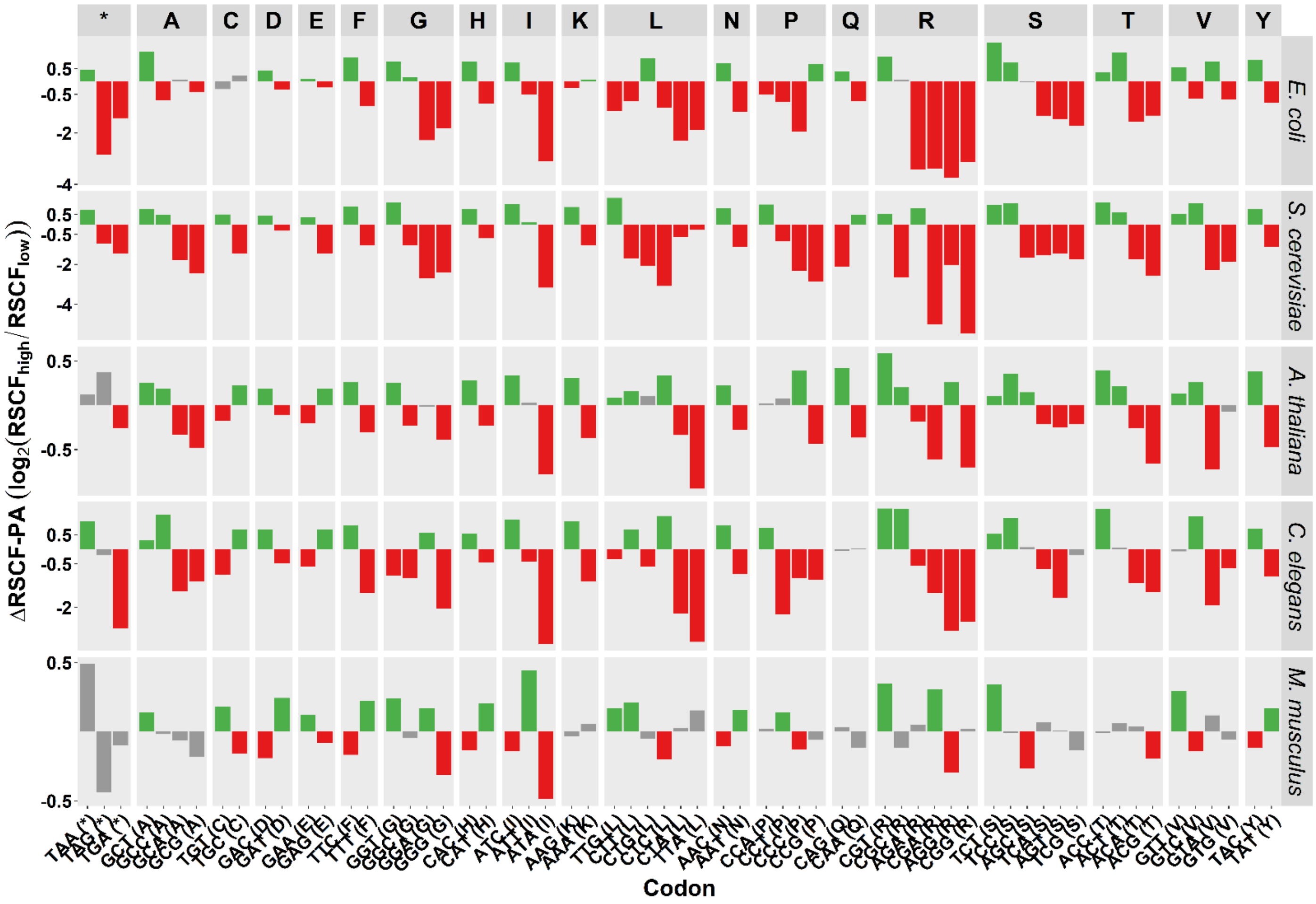

**Supplementary Figure S2.** Relative change in codon usage associated with protein abundance (ΔRSCF-PA). The ΔRSCF-PA values are calculated as the $\log_2$ ratio of the RSCF values of the highest abundant 5% ($RSCF_{high}$) and the lowest abundant 5% ($RSCF_{low}$). The colours indicate significant positive association with highly expressed genes (green) or negative association (red) (permutation, FDR < 0.05). The grey bars indicate non-significant associations.

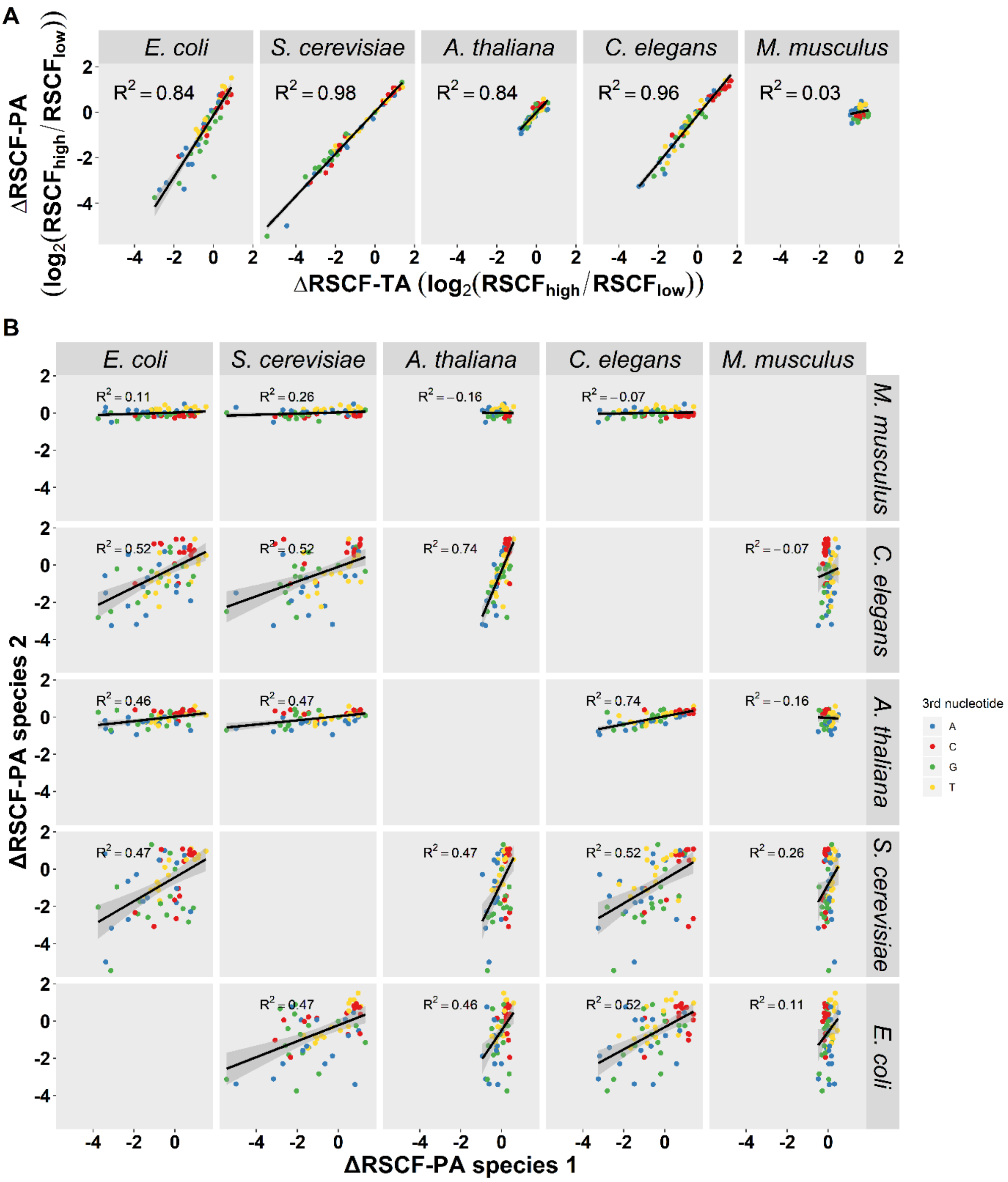

**Supplementary Figure S3**. Relations between relative changes in codon frequencies (ΔRSCF) associated with transcript and protein abundances. (A) The relation between ΔRSCF based on transcript abundance (ΔRSCF-TA) and protein abundance (ΔRSCF-PA) within species. The line plotted is based on a linear fit, for which the $R^2$ is indicated in the panel (Pearson correlation). (B) The relation between shifts in codon frequencies associated with protein abundance (ΔRSCF-PA) between species. As in supplementary figure 1, on the x-axis the ΔRSCF-PA values for species 1 are plotted (species name is indicated on the bar on top), on the y-axis the ΔRSCF-PA values for species 2 are plotted (species name is indicated on the bar on the right). The line is based on a linear model and added as visual aid, the reported correlation is the spearman correlation.

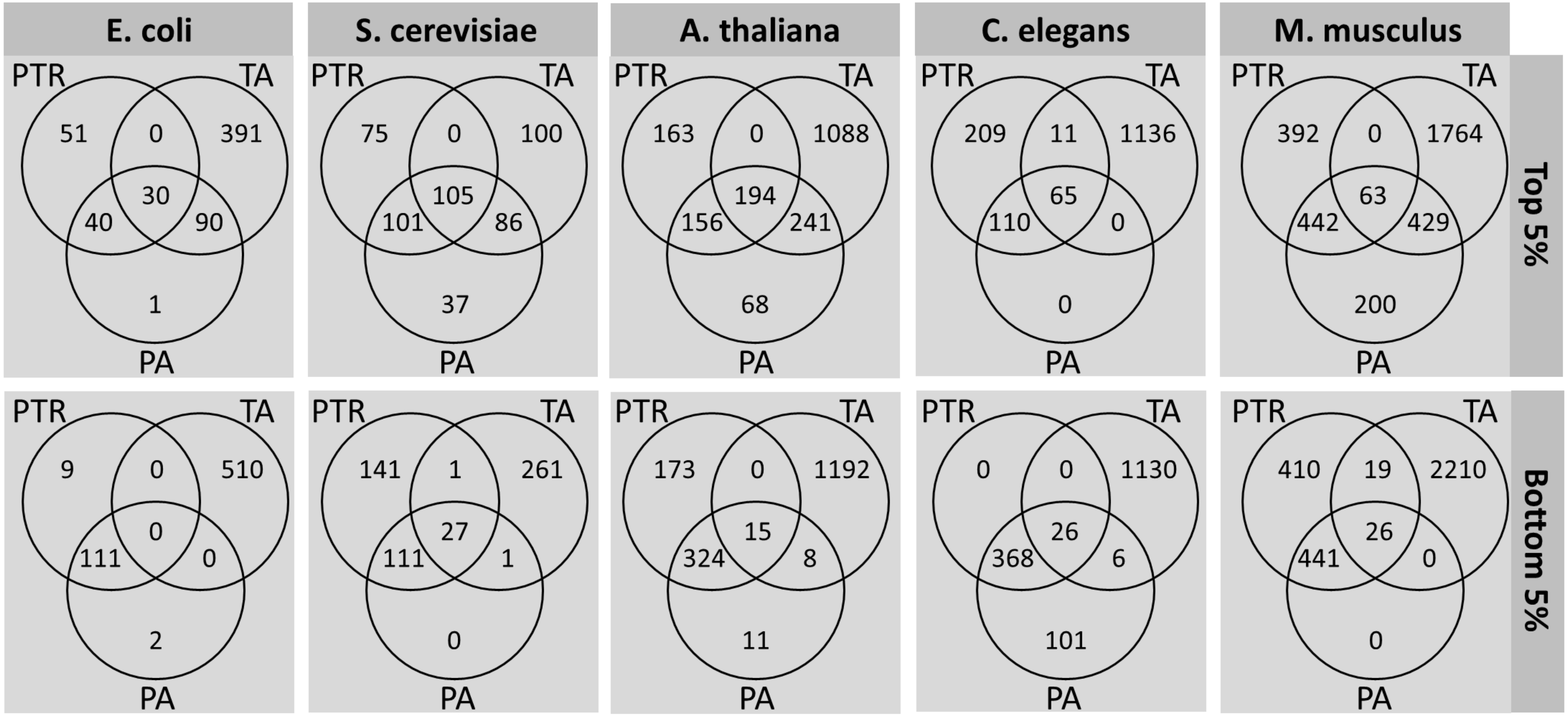

**Supplementary Figure S4**. Overlap of the gene pools per top and bottom 5%. Per species, split out for the top 5% and the bottom 5%, the overlap in genes belonging to the transcript abundance (TA), protein abundance (PA), and protein per transcript ratio (PTR) extremes.

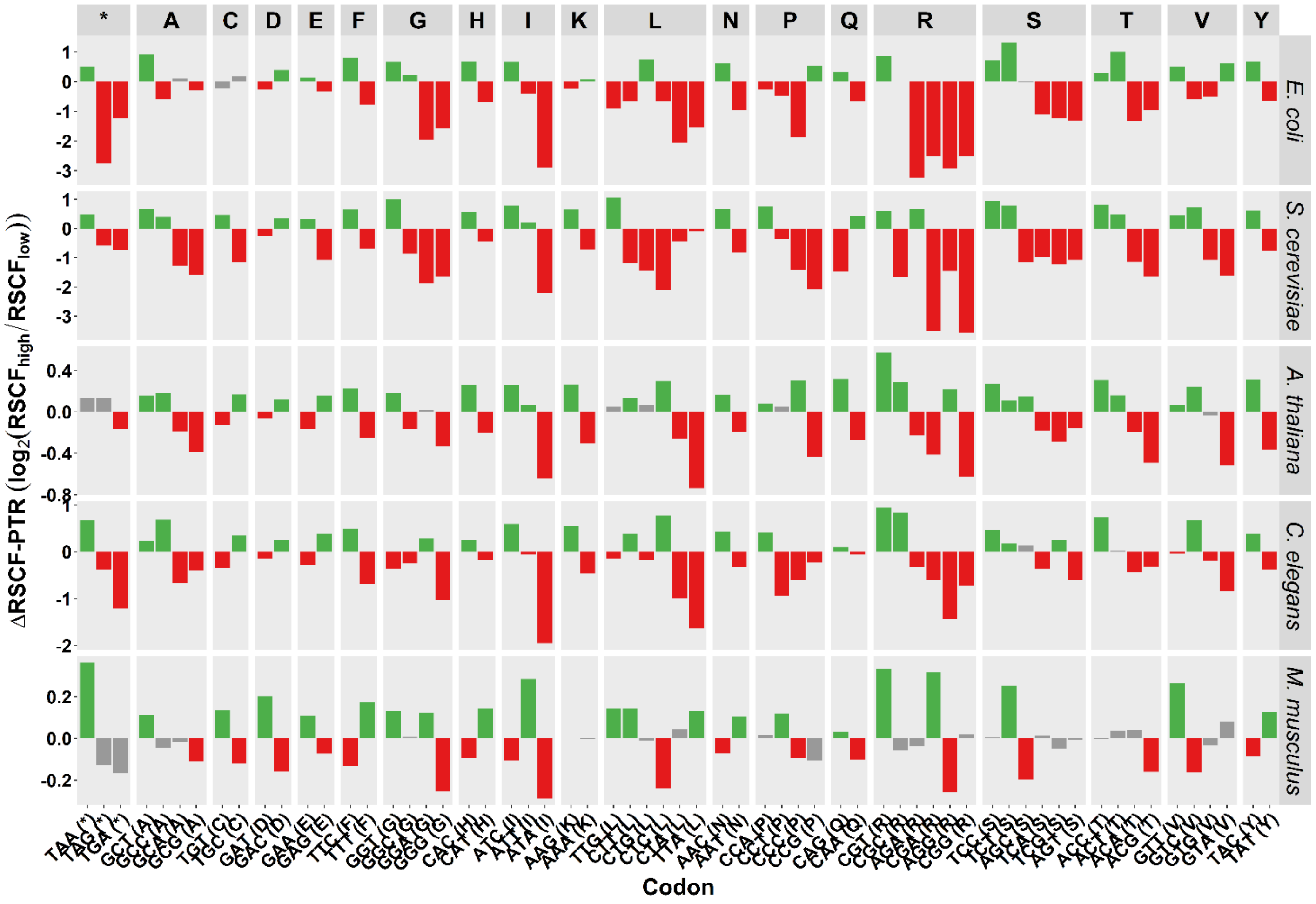

**Supplementary Figure S5**. Relative change in codon usage associated with protein per transcript ratio (ΔRSCF-PTR). The ΔRSCF-PTR values are calculated as the $\log_2$ ratio of RSCF values of the highest abundant 5% ($RSCF_{high}$) and the lowest abundant 5% ($RSCF_{low}$). The colours indicate significant positive association with highly expressed genes (green) or negative association (red) (permutation, FDR < 0.05). The grey bars indicate non-significant associations.

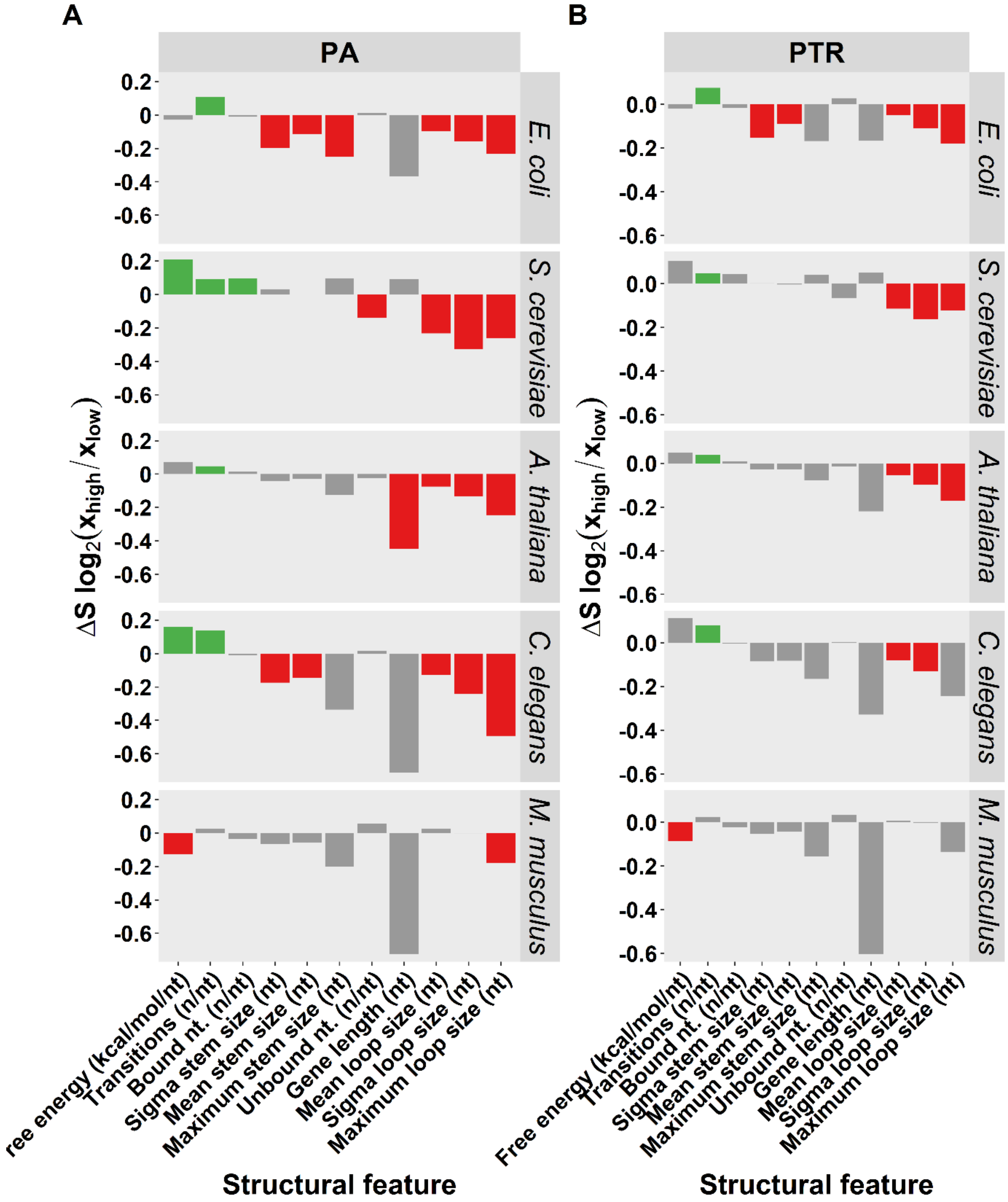

**Supplementary Figure S6.** Bias in secondary structure of mRNA associated with protein abundance (PA) and protein per transcript ratio (PTR). (A) Bias in structural features (ΔS) calculated as the $log_2$ ratio of the average structure value (e.g. number of nucleotides (nt) per maximum loop size) in the highest abundant 5% ($x_{high}$) and the lowest abundant 5% ($x_{low}$). Colours indicate if a particular feature is significantly positively associated (green) or negatively associated (red) with high transcript abundances (FDR < 0.05), grey bars indicate non-significant associations (FDR > 0.05). (B) As in A, but for the contrast between the 5% highest protein per transcript ratio versus the 5% lowest ratio.

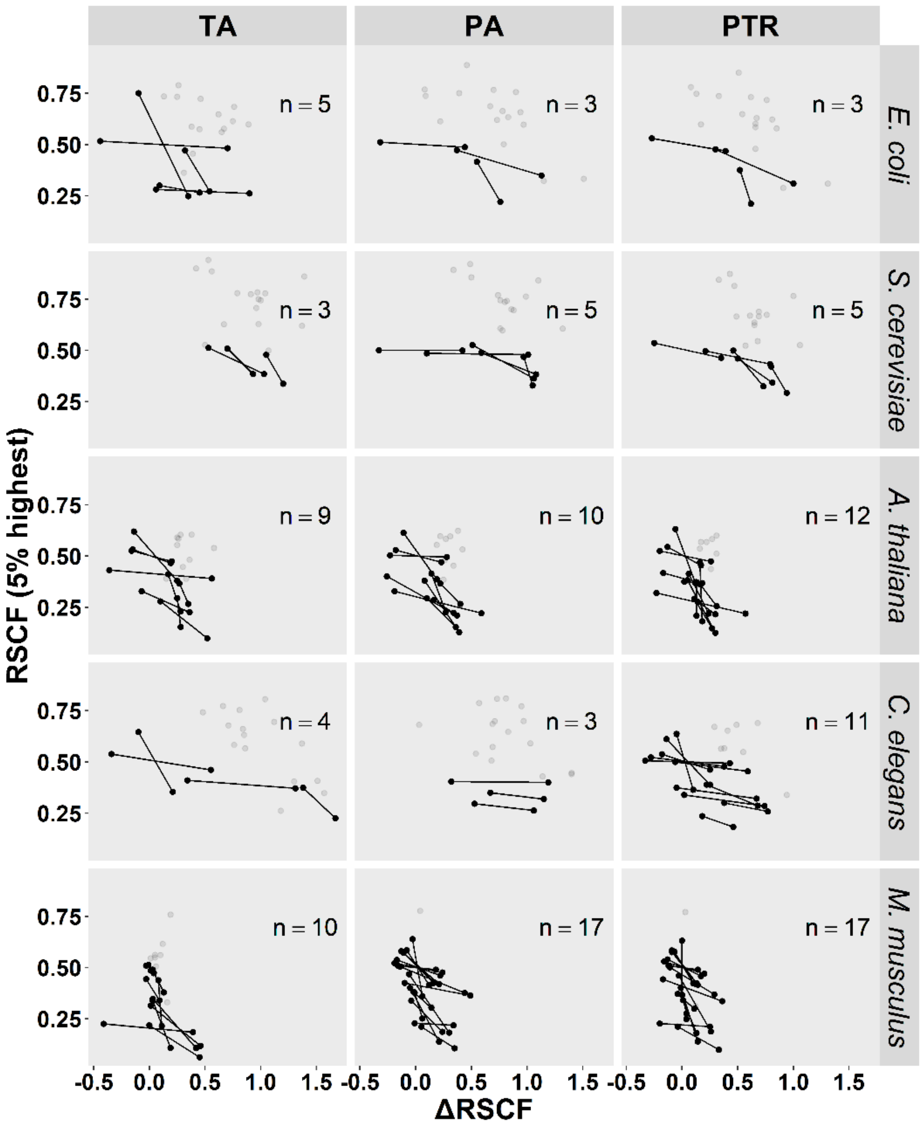

**Supplementary Figure S7**. Relation between codons showing the highest relative increase as determined by comparing the 5% lowest and 5% highest expressed genes (ΔRSCF; x-axis) and the most frequent in the top 5% highest expressed genes (RSCF; y-axis). This was split out per species (along y-axis) and type of ΔRSCF (along x-axis) based on the analyses of transcript abundance (TA), protein abundance (PA), and protein per transcript ratio (PTR). Each dot represents the synonymous codon that was associated with highest ΔRSCF and/or the highest relative synonymous frequency in highly expressed genes (RSCF). To clarify: if two connected dots are plotted, the most frequent codon in the top 5% highest expressed genes (RSCF) is not the same as the synonymous codon with the highest ΔRSCF value. The numbers given in each plotting window list how many of these codon-pairs existed (e.g. in *C. elegans* for three amino acids different synonymous codons were identified based on ΔRSCF-PA and RSCF -PA values)

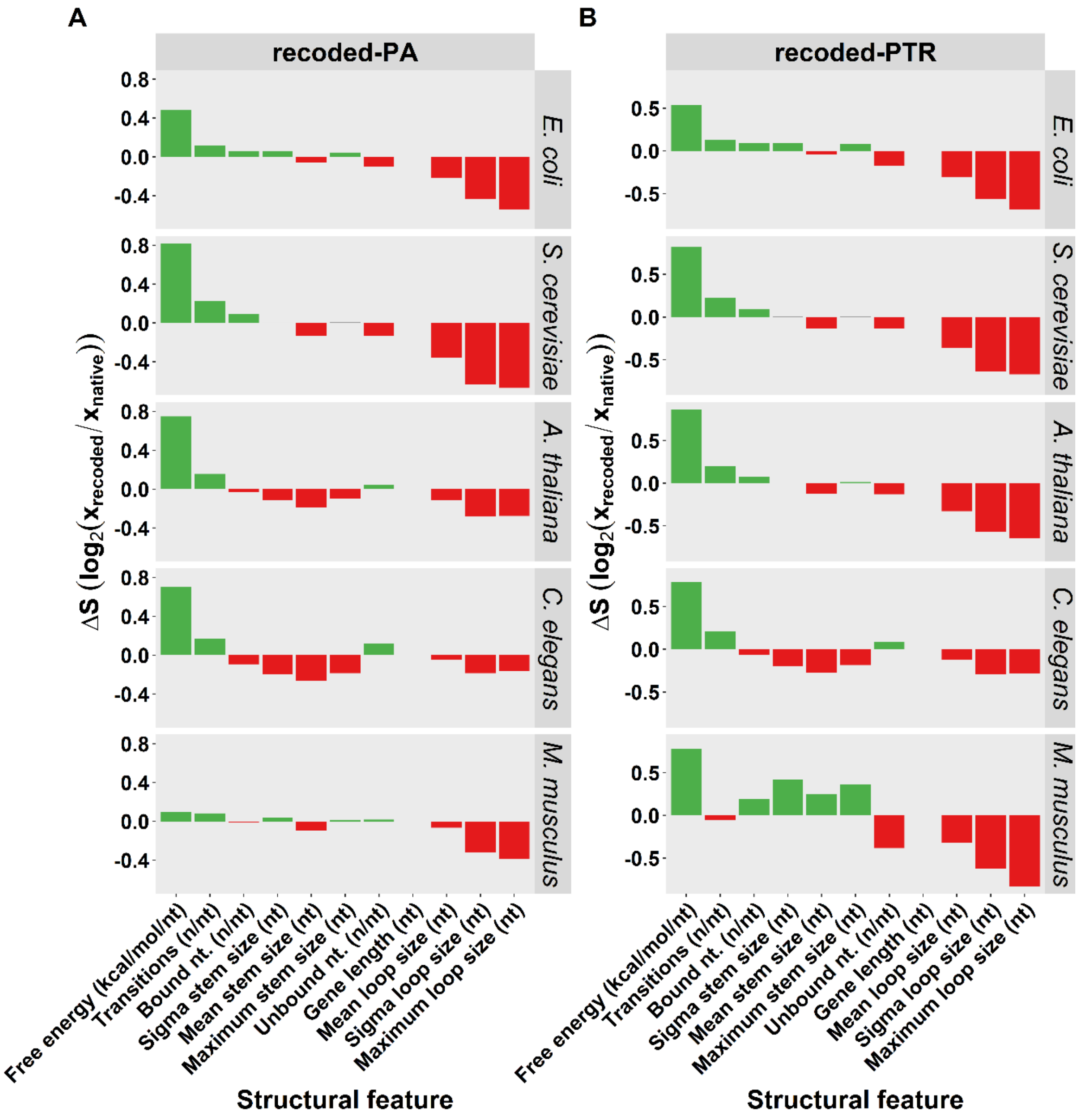

**Supplementary Figure S8.** Changes in mRNA secondary structures after whole-genome recoding. The codons for recoding were chosen by selecting for each amino acid a codon that increased most in frequency at high extremes as identified by analyses of protein abundances (recoded-PA) (**A**) and protein per transcript ratio (recoded-PTR) (**B**). Each recoded transcript was compared to its native counterpart. The shifts in structural features (ΔS) were calculated as the $\log_2$ ratio of the average structure value (e.g. number of nucleotides (nt) per maximum loop size) in recoded sequences ($x_{recoded}$) and native sequences ($x_{native}$). Colours indicate if a particular feature is significantly positively associated (green) or negatively associated (red) with high protein abundances (Bonferroni corrected paired t-test, $p < 0.05$), grey bars indicate non-significant associations (Bonferroni corrected paired t-test, $p > 0.05$).

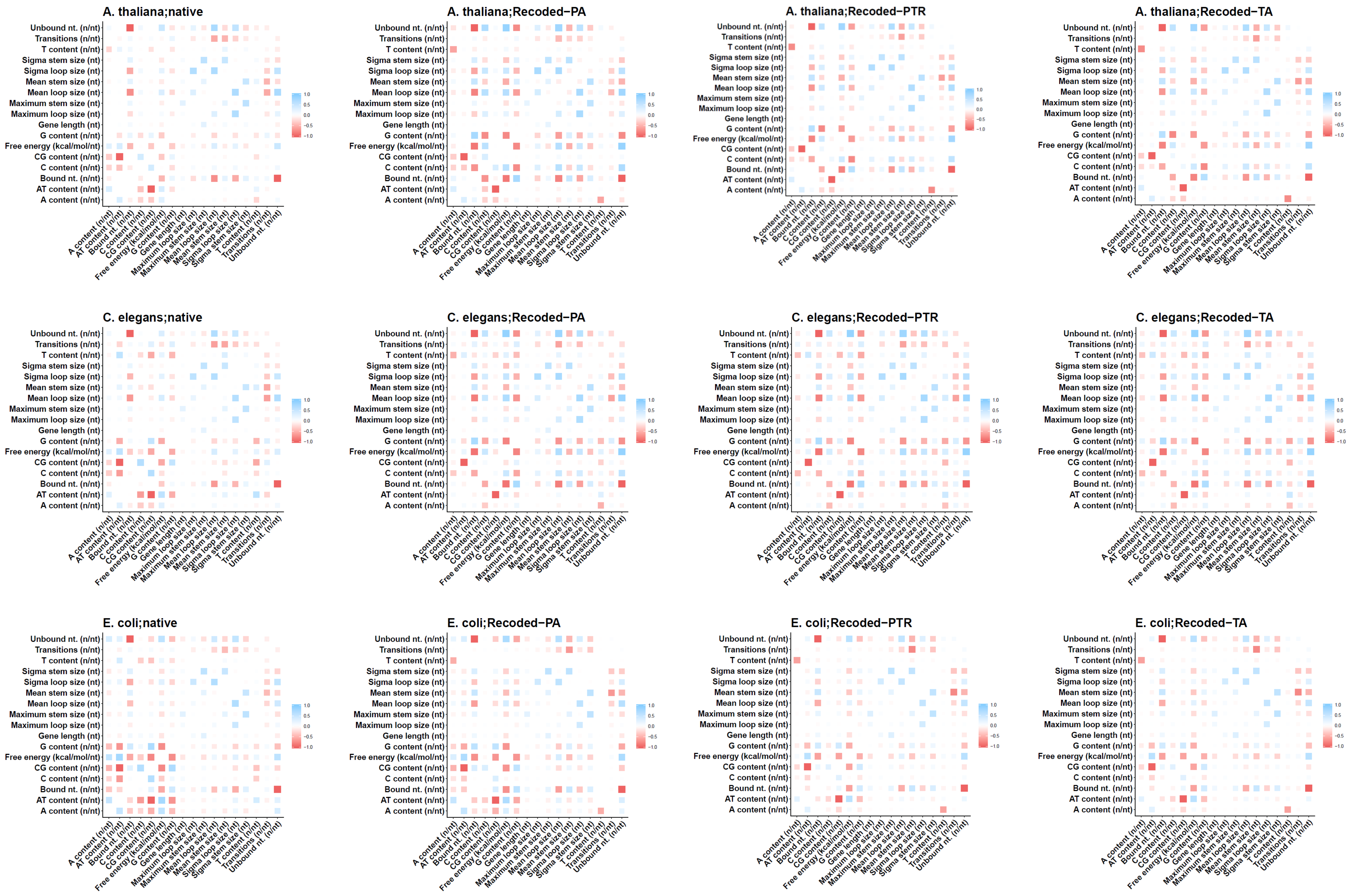

A. thaliana;native
A. thaliana;Recoded−PA
A. thaliana;Recoded−PTR
A. thaliana;Recoded−TA
C. elegans;native
C. elegans;Recoded−PA
C. elegans;Recoded−PTR
C. elegans;Recoded−TA
E. coli;native
E. coli;Recoded−PA
E. coli;Recoded−PTR
E. coli;Recoded−TA
Unbound nt. (n/nt)
Transitions (n/nt)
T content (n/nt)
Sigma stem size (nt)
Sigma loop size (nt)
Mean stem size (nt)
Mean loop size (nt)
Maximum stem size (nt)
Maximum loop size (nt)
Gene length (nt)
G content (n/nt)
Free energy (kcal/mol/nt)
CG content (n/nt)
C content (n/nt)
Bound nt. (n/nt)
AT content (n/nt)
A content (n/nt)
1.0
0.5
0.0
−0.5
−1.0

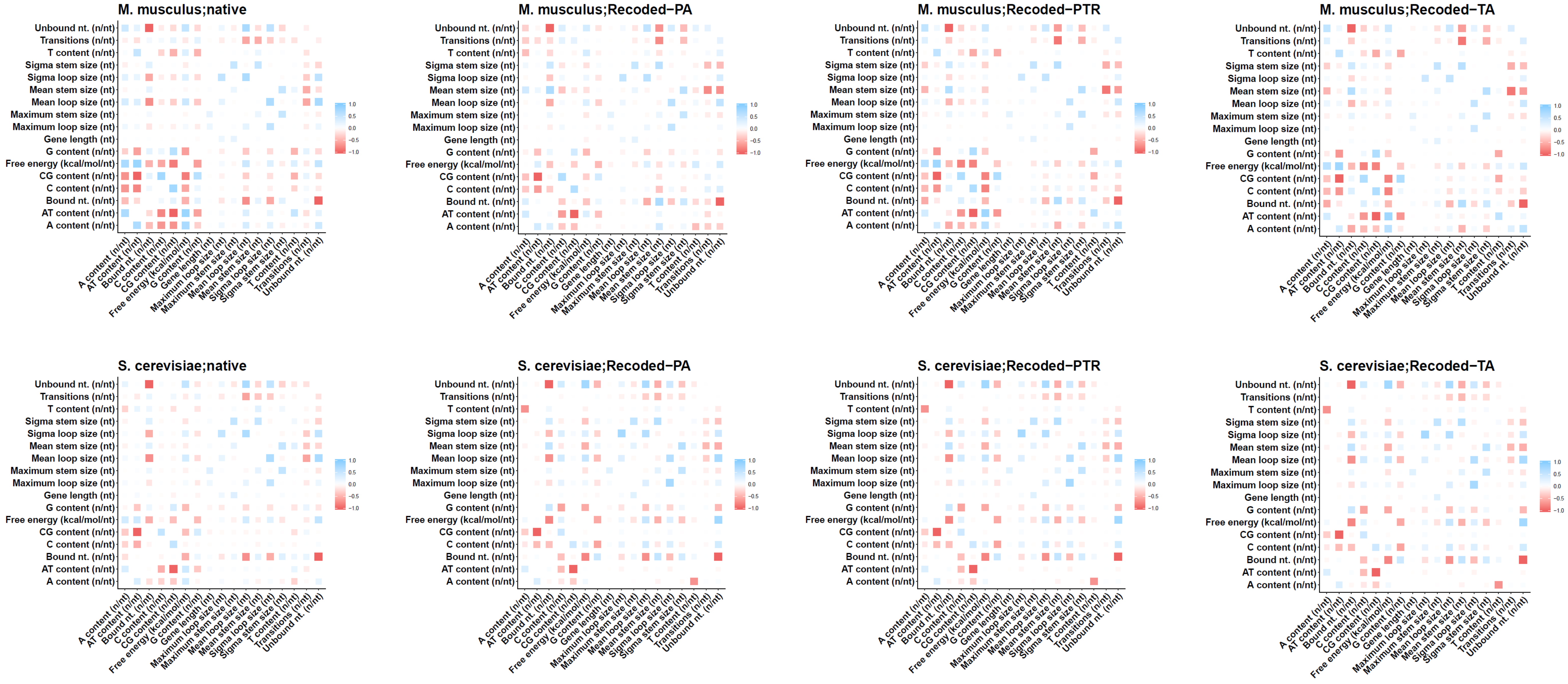

**Supplementary Figure S9**. Pearson correlation between structure parameters of mRNA secondary structures. These plots show the correlations between structure parameters for native and recoded mRNAs for all five species (*E. coli*, *A. thaliana*, *S. cerevisiae*, *C. elegans*, and *M. musculus*). For recoding of  mRNAs codons were selected that increased most in frequency at high extremes as identified by analyses of transcript abundances (Recoded-TA), protein abundances (Recoded-PA) and protein per transcript ratio (Recoded-PTR). Each plot shows the correlations within one set of mRNAs for a species. For a more summarized overview, see supplementary figure S10.

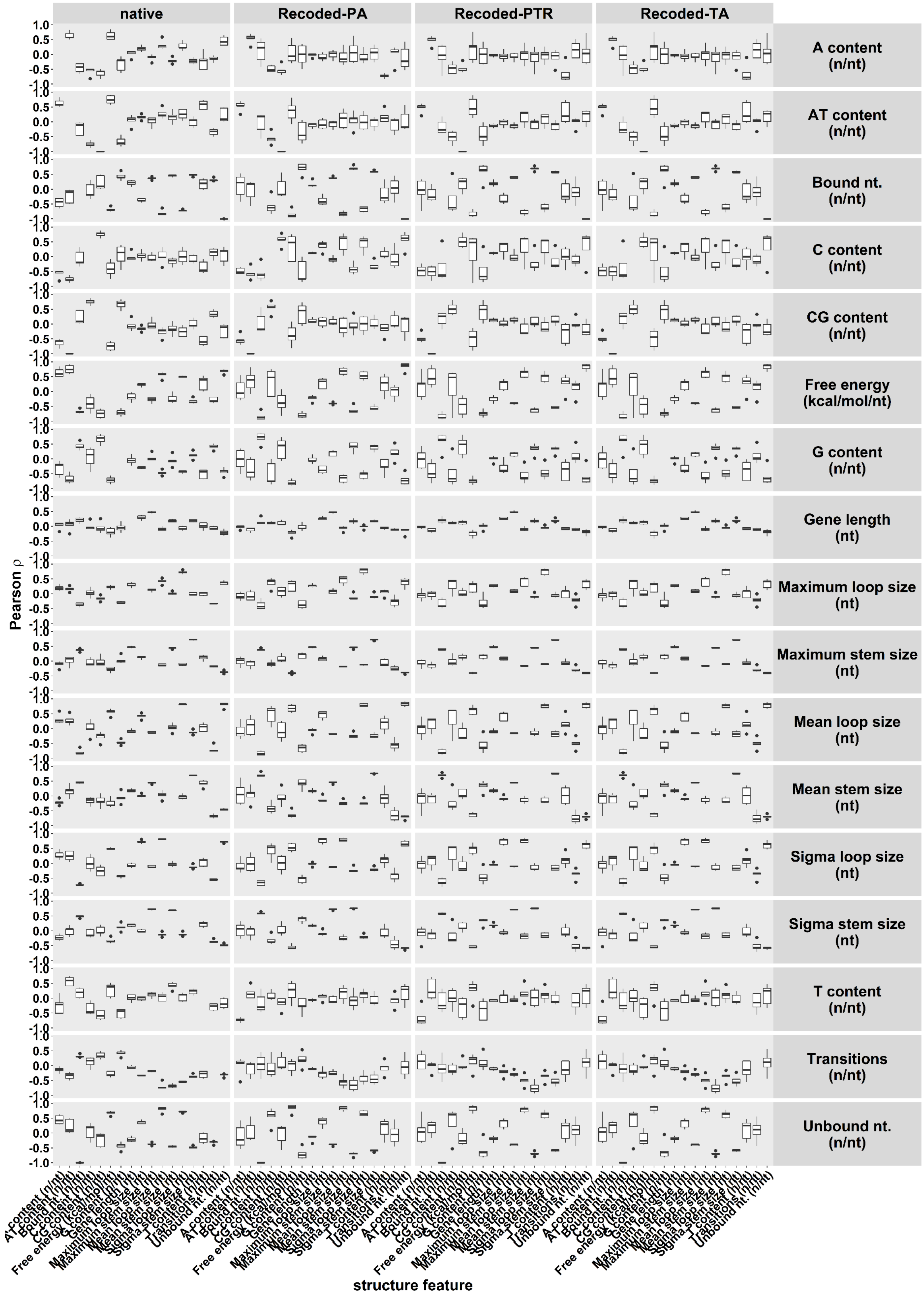

**Supplementary Figure S10.** Summarized overview of correlations between nucleotide content and structural features of native and recoded mRNA secondary structures. For recoding of mRNAs, codons were selected that increased most in frequency at high extremes as identified by analyses of transcript abundances (Recoded-TA), protein abundances (Recoded-PA) and protein per transcript ratio (Recoded-PTR). The x-axis shows the structural features, split out per type of mRNA (native, Recoded-PA, Recoded-PTR, and Recoded–TA), the y-axis shows the Pearson correlation coefficient. Each box-plot represent all five species (*E. coli*, *A. thaliana*, *S. cerevisiae*, *C. elegans*, and *M. musculus*). For details per species, see Supplementary Figure S9.
